# Measuring and monitoring skill learning in closed-loop myoelectric hand prostheses using speed-accuracy tradeoffs

**DOI:** 10.1101/2023.08.10.552753

**Authors:** Pranav Mamidanna, Shima Gholinezhad, Dario Farina, Jakob Lund Dideriksen, Strahinja Dosen

## Abstract

**Objective:** Closed-loop myoelectric prostheses, which combine supplementary sensory feedback and electromyography (EMG) based control, hold the potential to narrow the divide between natural and bionic hands. The use of these devices, however, requires dedicated training. Therefore, it is crucial to develop methods that quantify how users acquire skilled control over their prostheses to effectively monitor skill progression and inform the development of interfaces that optimize this process.

**Approach:** Building on theories of skill learning in human motor control, we measured speed-accuracy tradeoff functions (SAFs) to comprehensively characterize learning-induced changes in skill – as opposed to merely tracking changes in task success across training – facilitated by a closed-loop interface that combined proportional control and EMG feedback. Sixteen able-bodied and one amputee participated in a 3-day experiment where they were instructed to perform the box-and-blocks task using a timed force-matching paradigm at four specified speeds to reach two target force levels, such that the SAF could be determined.

**Main results:** We found that the participants’ accuracy increased in a similar way across all speeds we tested. Consequently, the shape of the SAF remained similar across days, at both force levels. Further, we observed that EMG feedback enabled participants to improve their motor execution in terms of reduced trial-by-trial variability, a hallmark of skilled behavior. We then fit a power law model of the SAF, and demonstrated how the model parameters could be used to identify and monitor changes in skill.

**Significance:** We comprehensively characterized how an EMG feedback interface enabled skill acquisition, both at the level of task performance and movement execution. More generally, we believe that the proposed methods are effective for measuring and monitoring user skill progression in closed-loop prosthesis control.

## Introduction

Myoelectric prostheses seek to restore the lost manipulation capabilities of individuals with a limb difference by using electromyographic (EMG) signals to estimate the user’s motion intent to drive powered prosthetic devices. Closed-loop user-prosthesis interfaces, which combine EMG-based control and artificial sensory feedback to provide users with a sense of prosthesis state and improve the intuitiveness of control, hold the promise of bridging the gap between natural and bionic hands [1]–[3].

To use these sophisticated devices and interfaces, the prospective user needs to train to perform specific, often unintuitive, contractions of their residual muscles. Attaining expert level skill in user-prosthesis interaction may require guided training over weeks and months [4]–[6]. This relatively long learning process is due to the inherent variability of EMG-based control and non-linear properties of the prosthesis, such as non-back-drivability. Quality training is crucial for successful control and poor training has been linked to device rejection [7]. Training is also needed for the feedback side of the interface since the user needs to learn to perceive the sensations artificially elicited [2] and to use this sensation to gain information on the prosthesis state. Quantifying the way and time course of the acquisition of skilled prosthesis control is a fundamental step to develop evidence-based training or rehabilitation protocols [8], [9], design interfaces that are easier to learn and retain, and understand the motor learning processes underlying complex human behaviors [10].

Previous studies that investigated prosthesis skill learning, have defined it in terms of improved accuracy, faster movement speed, reduced end-point variability or a combination thereof [8], [9], [11]–[13]. These studies mainly evaluated skill by asking participants to be as accurate as possible in the given task at a self-chosen comfortable pace, or as fast as possible. However, this approach neglects the effect of the self-chosen speed on the resulting movements and performance. Movement speed and accuracy are fundamentally coupled, and skill can only be inferred when both change in the expected direction [14]–[18]. Figure 1A illustrates this concept with a simple example. If the speed-accuracy measurements over two days for a participant is given by the filled circles as shown in the figure, it would be impossible to dissociate if the improved accuracy (blue circle) is due to better skill or merely due to performing the task at a slower speed (as compared to the gray circle). Therefore, determining speed-accuracy trade-off functions (SAFs, Figure 1A) has been proposed as a solution to rigorously study skill learning [14], [16], [19].

**Figure 1:**
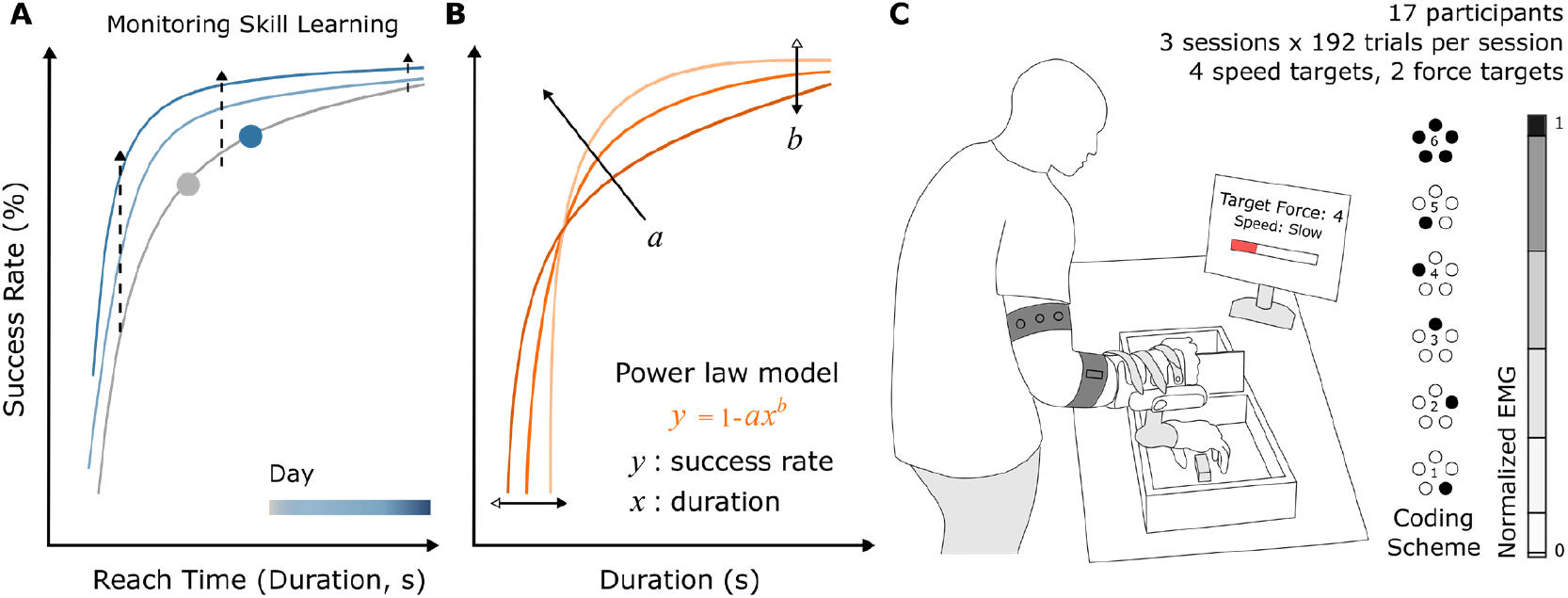
Monitoring skill by measuring and modelling learning induced changes in SAF – concept and experimental setup. (A) Each curve represents the SAF resulting from the use of a closed-loop prosthesis interface, over a period of training. Plausible changes to the SAF across training are indicated by dashed arrows. (B) Diagram showing the power law model of SAF and the effect of parameters *a* and *b* on it. The model was used to capture the changes in SAF observed experimentally. Changes in model parameters can be used to quantify the (subject-specific) differences in skill across time. (C) Experimental setup shows an able-bodied participant using a closed-loop prosthesis interface, combining dual-site proportional control and vibrotactile EMG feedback, to control the Michelangelo hand prosthesis. Salient experimental protocol details are also shown in the top right; EMG feedback spatial coding scheme is displayed on the right. Modified with permission from [22].

This approach evaluates task performance systematically and comprehensively by asking the participants to repeat the same task at different speeds. A SAF so measured, characterized by its shape (monotonically increasing), rate, and intercept, succinctly captures skill [20], [21]. Previous work in human motor control demonstrated that SAF changes across days in a way that is consistent across speeds [14], [19], and is mainly driven by better motor execution, reflected by decreased movement variability. However, such a detailed analysis is lacking for prosthesis control – it is unknown if SAF changes such that improvements are similar across speeds, or if some speeds experience larger gains than others (e.g., faster speeds might have larger gains as opposed to slower speeds, simply by having more scope for such gains (Figure 1A, dashed arrows)).

Apart from quantifying skill, SAFs have been used in a recent study to understand and compare how different feedback interfaces afford different speed-accuracy tradeoffs in prosthesis force control [22]. The study demonstrated the relevance of SAFs and showed how comparing the interfaces at a single speed (corresponding to single point on the SAF profile), as commonly done in the literature, could generate misleading conclusions. However, it is still unknown how the tradeoff evolves across days during learning, or in other words, how skill develops over time. Quantifying learning-induced changes in the SAF enables us to thoroughly investigate how users acquire skills, and accordingly develop tools to monitor and track skill acquisition and retention.

In relation to the latter, Reis et al., [14] proposed a model-based method to monitor skill progression. Given that the SAF reflects skill, it follows that if a parametric model of SAF can be obtained, then changes in the parameter values of such a model could be used to monitor skill over time, in a subject specific manner (Figure 1B). Notably, they proposed a simple 1-parameter model of SAF such that the single parameter reflected changes in skill. Similarly, Guiard et al., [23] observed that the SAF in a cursor pointing task could be captured using a 1-parameter power law model. However, it remains to be seen if such methods generalize to skill monitoring in prosthesis control.

In this study we therefore characterize the SAF resulting from the use of a closed-loop user-prosthesis interface for grip force control, to investigate skill learning and the behavioral changes subserving it. Participants used a closed-loop interface that included a dual-site direct proportional control of prosthesis closing speed and EMG-feedback [24], [25]. In this approach, an array of vibrotactors was employed to deliver the level of the myoelectric signal (prosthesis command input) as the feedback to the participants, thereby facilitating predictive control of grasping force by augmenting a sense of contraction strength. Over the course of 3 days, participants practiced a modified version of the box-and-blocks task where the blocks were required to be grasped at specified forces and speeds (timed force-matching paradigm). We hypothesized that the largest gains in accuracy would be experienced at faster speeds, due to a larger margin for improvement (as explained above) as well as the predictive nature of the feedback. We then built a parametric power-law model of the measured SAFs (Figure 1B), following previous studies in motor skill learning [14] and human-computer interaction [23], [26]. Using the model, we characterized how the SAF changed across days for this type of closed-loop control interface and compared the results to those reported in the literature for human motor control of natural movements. Finally, we discuss if skill could be inferred from “observational” data, where participants are simply asked to perform the task at their self-chosen comfortable pace to estimate the parameters of the SAF, such that the process of measuring the true SAF could be simplified.

## Methods

### Participants

Sixteen healthy, able-bodied participants (nine males and seven females, aged 23.5 ± 2.6 years) were recruited. One transradial amputee (female, 50 years old, 11 years since traumatic amputation of non-dominant hand, limited use of a single degree of freedom (DoF) myoelectric prosthesis), who had prior experience with similar experimental setups was also recruited for the study. All participants signed an informed consent form before the start of the experiment. The experimental protocol was approved by the Research Ethics Committee of Region Nordjylland (approval number N-20190036).

### Experimental Setup

The experimental paradigm follows from a previous experiment [22], which for the present study was extended across days. The methods are briefly described here. The experimental setup is shown in Figure 1C. Participants wore an orthotic wrist immobilization splint to achieve near-isometric wrist flexion and extension movements. The Michelangelo prosthetic hand (OttoBock GmBH, DE) was attached to the splint, with the arm placed in a neutral position. Further, two dry EMG electrodes with embedded amplifiers (13E200, Otto Bock, DE) were placed over the wrist flexors and extensors of the right forearm. The electrode placement was determined by visually observing muscle contractions and palpating the forearm. Five vibrotactors (C-2, Engineering Acoustics Inc.) were positioned evenly around a cross-section of the upper arm,and secured in place by an elastic band. A modified box-and-blocks setup was used for the experimental task, where the participants were asked to apply predefined forces when grasping the blocks and to perform the task within a set time interval, as explained in the section “Experimental Design”. The prosthesis was connected to a standard laptop PC via Bluetooth, while the vibrotactors were connected via USB. The task instructions were displayed on a dedicated computer screen, positioned at a comfortable viewing angle and distance. The experiment was developed in MATLAB Simulink using a toolbox for testing human-in-the-loop control [27]. The control loop operated in real-time on the host PC at 100 Hz, through the Simulink Desktop Real-Time toolbox. In the case of amputee participant, the prosthesis was attached to a custom-made socket with integrated EMG electrodes, while the vibrotactors were placed on the upper arm, using the same setup as for the able-bodied participants.

### EMG Control

Participants used near-isometric wrist flexion and proportional control to generate velocity commands to close the prosthesis. However, since fine control of prosthesis opening was not relevant for the study, it was simply triggered by a strong contraction of the wrist extensors [22]. The linear envelope of EMG was sampled at 100 Hz from two electrode systems, placed on the flexors and extensors, as explained above, through the embedded prosthesis controller and transmitted to the host PC. The signals were subsequently filtered using a 2^nd^ order Butterworth low-pass filter with a 0.5 Hz cutoff, and this ‘myoelectric command’ from each of the electrodes was normalized to 50% of that obtained during maximum voluntary contraction (MVC) (following the results of [28]). The normalized myoelectric commands obtained from the flexor EMG were translated into the normalized input command for the prosthesis using a piecewise linear mapping defined by the breakpoints given in Table 1 (see [22]). Note that the EMG intervals were wider for higher amplitudes to compensate for the higher variability of the EMG at stronger contractions. The prosthesis input command determined closing velocity (1 – maximum velocity) and generated grasping force, i.e., the stronger the command input, the faster the prosthesis closes and the higher the force generated after contact. The aforementioned mapping effectively translated the EMG “levels” into the corresponding force levels (see the next section for a more detailed explanation of closed-loop control). Note that for EMG amplitude lower than 0.025 on the normalized scale, no input command was sent to the prosthesis (Level 0) to avoid unintentional prosthesis closing. To trigger prosthetic hand opening, however, participants simply needed to reach 0.4 on the normalized range (i.e., 20% MVC) of extensor EMG.

**Table 1:** The myoelectric command, prosthesis input and force ‘level’ boundaries for the discretized interface.

| Level | 0 | 1 | 2 | 3 | 4 | 5 | 6 |
| --- | --- | --- | --- | --- | --- | --- | --- |
| Myoelectric command | 0.025 | 0.1 | 0.27 | 0.47 | 0.69 | 0.95 | 1 |
| Prosthesis input | 0 | 0.25 | 0.42 | 0.59 | 0.76 | 0.9 | 1 |
| Force | 0.05 | 0.31 | 0.45 | 0.58 | 0.73 | 0.9 | 1 |

### Vibrotactile EMG Feedback

The flexor EMG command generated by the participant was delivered as feedback through a vibrotactor array. A spatial coding scheme consisting of six stimulation patterns was used to convey the six discrete levels of the myoelectric command. To represent the first five levels, one of the tactors from the array was activated individually in a specific order. However, the sixth level was indicated by activating all the tactors simultaneously (see Figure 1C). During setup, adjustments were made to the position and/or intensity of the vibrotactors if they caused unpleasant or poorly localized sensations, to ensure that the participants could easily differentiate all the stimulation patterns. The vibration frequency for all tactors was set to 200 Hz, and the stimulation pattern was updated at 50 Hz.

The six discrete levels were defined using the same piece-wise linear mapping reported in Table 1. As soon as the participants contracted their wrist flexors (to generate a myoelectric command above level 0), they received feedback about the level (1-6) they were generating, thereby enabling them to modulate the prosthesis command input to the desired level using online feedback. The control and feedback interfaces were designed in such a way that if the participants generated a particular level of myoelectric command, the prosthesis applied the same level of force on the object. For instance, if a participant generated and maintained commands corresponding to level 2, the prosthesis closed at level 2 velocity and exerted level 2 force onto the grasped object. Note that while the feedback provided discrete information (a level of EMG), the command signal generated by the participant and transmitted to the prosthesis was still continuous and proportional (as in commercial approach to prosthesis control). Therefore, if the EMG signal was in the higher range of level 2 (e.g., close to the threshold between level 2 and 3), the generated grasping force would be in the higher range of level 2 of prosthesis forces, as defined by Table 1.

### Experimental Design

The experiment was designed as a longitudinal trial over three sessions with a maximum of two days between the sessions. In each session, the participants trained to perform the box-and-blocks task with the timed force-matching requirements. In each trial, they were required to (1) apply a specified level of force on the object (either level 3 or 5, see Table 1), and (2) reach the target level within the specific time. Participants were required to perform the task in four speed conditions – Slow, Medium, Fast and Very Fast – which corresponded to time of execution in the range 4 – 8s, 3 – 5s, 2 – 4s, and 1 – 3s, respectively. The times of execution were defined to capture the relevant domains of the SAF curve and expand upon previous investigations [22]. Thereby, we used a time-band methodology to measure the SAF [20], [22], and with respect to our previous work [22], one more speed was introduced in the protocol in order to better sample the shape of the SAF profile. In addition to performing trials with both speed and force targets, the amputee participant also performed ‘observational trials’ where she was instructed to perform the same task at a self-chosen comfortable pace, with just the force target specified.

### Experimental Protocol

Initially, all equipment (the wrist immobilization splint, prosthesis, EMG electrodes and vibrotactors) were placed on the participant. Then, a brief calibration and familiarization followed in all sessions. During EMG calibration, 3 repetitions of 5-s long MVC contractions were recorded independently for both the flexors and extensors. The MVC measurements were recorded in the same posture that the participants would use to perform the box-and-blocks task, to account for the effect of arm posture and prosthesis weight on the recorded EMG. Next, the participants were guided to explore the control interface, i.e., the relation between muscle contraction intensity and velocity in hand opening/closing. Subsequently, they were familiarized with the vibrotactile feedback interface. This involved a spatial discrimination task where they were presented with two sets of 18 stimulation patterns (3 repetitions x 6 levels, Figure 1C) and asked to identify the patterns. The experiment proceeded once the participants achieved at least 95% success in the discrimination task, which usually required less than 5 min since the vibration patterns were designed to be easy to discriminate.

After familiarization with the control and feedback interfaces, participants performed 6 blocks of 32 trials (8 trials × 4 speed conditions) of the timed force-matching task. On the first day, an extra (familiarization) block of 32 trials was performed where the participant was guided through the task. In each trial, the force and speed targets were first displayed. Then, the participants used the proportional control interface and the provided EMG feedback to successfully complete the trial, as explained in section ‘Vibrotactile EMG feedback’. Once the participant felt they successfully reached (or overshot) the target, they were instructed to trigger prosthesis opening. As the prosthesis was non-backdrivable, like most commercial devices, it was not possible to decrease the grasping force in a controlled manner (the prosthesis would open and release the object completely). Therefore, as explained before, the overshoot was also considered as a reason to open the hand and end the trial. Immediately after the trial ended, they received feedback on their performance, in terms of both target force and target speed. During the familiarization trials, the participants were instructed on how to modulate their muscle contraction to control the closing velocity of the prosthesis. In each block of 32 trials, the target speed (Slow, Medium, Fast or Very Fast) was presented in a blocked fashion, i.e., it remained unchanged for 8 trials within which the target forces (3, 5) were presented 4 times each in a random order. The target speeds were presented in a balanced random order, across blocks.

The experimental protocol for the amputee case study was slightly modified, where she performed 5 blocks (instead of 6) of trials with both force and speed targets on all three days and performed 2 additional blocks of the same task but with no speed target (referred to as ‘observational trials’) on Day 2 and Day 3.

### Outcome Measures

During each trial, the recorded myoelectric commands and force measurements were processed to obtain the primary outcome measures – reach time and trial success, respectively. Reach time was calculated as the duration between the time when participant started generating the EMG input (above Level 0) and when the maximum force was reached during the trial. Therefore, a successful trial was defined as one where the reach time satisfied the speed requirement, and the maximum reached force was within the corresponding target force level. The trials were aggregated per speed condition to obtain percent success rates.

Further, to understand how the participants planned and executed the task across the sessions, we computed three behavioral metrics [22], [29]. We computed (1) the number of force corrections per trial that the participants made, by counting the number of distinct plateaus (longer than 250 ms) in the force trajectory [22], to gain insights into their control policies – specifically corresponding to how they included the supplementary feedback in their commands. Since EMG feedback promotes predictive modulation of commands, by allowing participants to adjust their muscle contraction during prosthesis closing so that the target force is generated at once, upon contact, without requiring further modulations [11], [25], the more successful the participants are in using this strategy, the lower the number of corrections. Further, we analyzed the quality of generated EMG commands, to understand if participants generated (2) smoother and (3) more repeatable EMG commands across days and speed targets. To evaluate smoothness, we calculated the spectral arc length (SPARC) of the EMG commands [30]. We chose SPARC as an alternative to jerk-based measures, because it is a more general measurement of smoothness of any movement related variable and is robust to high-frequency changes in the recorded signal. Notice that the SPARC values are negative, with higher values indicating smoother trajectories. To measure the repeatability of commands, we computed the trial-by-trial variability as follows: we first normalized all commands to 200 time points between the start of the trial and reach time, and then measured the standard deviation at each of the 200 time points. As the final measure of variability, we computed the median of the obtained standard deviations across the time points.

### Parametric Models of SAF

To capture the changes in the shape of the SAF curve across training (see Figure 1B), we built a parametric model of SAF as *y* = *f*(*x*; *a, b*), where *y* is success rate, *x* is reach time (duration of trial), and *f* is a function of *x* and two parameters *a, b* bearing functional forms already investigated in the literature. Specifically, we considered the power law equation described in [23], [26], relating the success rates and reach time developed in the context of 1-d cursor movements. While different in the functional form, the power law model is similar to (and simpler than) another model developed in the motor learning literature for a task that involved the production of a sequence of isometric finger forces [14]. Here we modified the equation to reflect the x-intercept (or asymptote) at 0.8 s, as it is impossible to get a successful trial below 0.8 s, due to limitations in the speed of the prosthesis response and EMG command generation:

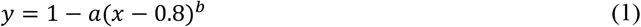

Note that parameter *b* in equation (1) is restricted to be negative.

### Model Fitting

The models were fit for day-wise pooled able-bodied participants’ data (success rate and reach time) for each of the 3 days, using ordinary least squares-based fitting. Success rates and reach times from 2 consecutive blocks were averaged so that the mean success rate and its variance are well represented during model fitting. This way, the power-law model was fitted using 12 points (4 speeds x 3 points per speed). The parameters obtained through this fitting procedure are denoted by asterisks, such as (*a*^*^, *b*^*^). The goodness of fit was measured using the R^2^ metric.

All models were log-transformed before fitting so that the parameters obtained would be easy to interpret. This way, parameter ‘log *a*’ denotes the y-intercept of the SAF in the log-log plane, while ‘*b*’ denotes the slope at which the success rate increases with reach time. A change (exclusively) in the intercept therefore indicates that the log-success rates at all movement speeds are affected similarly (e.g., consistent increase at all speeds), while a change in slope indicates that different speeds may affect success rate differently (e.g., increase at low speeds and decrease for high speeds).

### Reducing model complexity: 2-parameter to 1-parameter models

Earlier studies have observed that the SAF models described above can be further simplified such that only a single parameter reflects skill related changes [14], [23], but as explained before, this has been established for motor control tasks involving natural movements. Here, in a similar vein, we investigated if one parameter reflected skill improvements while the other parameter could be fixed in the context of closed-loop prosthesis control. In other words, we were interested to find if *f*(*x*; *a, b*) ≈ *f*(*x*; *a, <u>b</u>*) or *f*(*x*; *a, b*) ≈ *f*(*x*; *<u>a, b</u>*), where the underscore denotes the parameter being held constant. To start, let us denote the R^2^ obtained by fitting a particular set of data {*y, x*}_*i*_ to the 2-parameter model such that 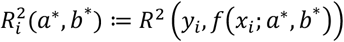, where *i* = {1, 2, 3} denotes the pooled success rate (*y*_*i*_) and reach time (*x*_*i*_) dataset of a particular day and *a*^*^, *b*^*^, denote the best fit parameters.

Then, Δ*R*^*2*^ represents the change in the goodness of fit (R^2^) when using the 1-parameter model, e.g., obtained by fixing *a* constant at *a*^*^ and setting the *b* parameter to *b*_*k*_ in the vicinity of *b*^*^. A significant decrease in R^2^ due to such a perturbation in one of the parameters (*a* or *b*) indicates that perturbing that parameter leads to worse fits with respect to the full 2-parameter model. By quantifying the sensitivity of the model fits to both parameters, we can rigorously quantify how they affect the goodness of fit and choose which parameter to set as a constant. Accordingly, we selected 1000 values of *a* and *b* in the range of the parameters in the vicinity of those observed by fitting the experimental data, (*a*_*min*_, *a*_*max*_) and (*b*_*min*_, *b*_*max*_) and computed the following set of R^2^ values:

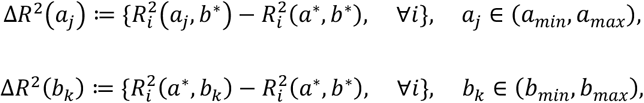

where index *i* = {1, 2, 3} denotes the *day*. We then obtain the simplified 1-parameter model by finding the parameter that changed mean R^2^ the least,

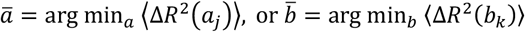

where ⟨. ⟩ denotes the mean of the set of values Δ*R*^*2*^.

### Statistical Analysis

3-factor repeated measures ANOVAs for success rate and behavioral measures were fit with 3 within-subjects factors – day, target speed and target force. To analyze the total improvement between Day 1 and 3, we fit 2-factor ANOVAs with target force and target speed as the factors. The assumptions of normality, homogeneity of variance and sphericity were verified using Shapiro-Wilk’s, Levene’s and Mauchly’s tests, respectively. Post-hoc analyses for all quantities of interest were performed by using pairwise t-tests, adjusted using the Holm-Bonferroni method. The threshold for statistical significance was set at p < 0.05. Mean ± standard deviation of outcomes per group of interest are reported throughout the paper.

## Results

### Learning induced changes in the SAF

We analyzed how participants’ SAFs (see Figure 2A) changed across the experiment by fitting a 3-way ANOVA with day, target speed and target force as factors. We found that all three factors significantly influenced the success rate (day: F(2, 30)=14.05, p=5e-5, target speed: F(3, 45)=35.47, p=6e-12, and target force: F(1, 15)=9.46, p=7e-3), but none of the factor interactions was significant. Participants exhibited a wide range of success rates across days and speeds, ranging from 66.3 ± 15.2% in the Very Fast speed on Day 1, to 90.2 ± 10% in the Slow speed on Day 3. Post-hoc analyses revealed that success rates improved significantly from Day 1 to 2 (p-adj=0.01), while the difference from Day 2 to 3 was not significant (Day 1: 75.4 ± 14.6%, Day2: 81.7 ± 13.3%, and Day 3: 85.3 ± 10.8%; average success rates across speeds). In a similar way, the participants’ success rates significantly decreased from Very Fast to Fast (p-adj=0.005) and Fast to Medium (p-adj=7e-4) but not from Medium to Slow speed. A significant effect of target force meant that participants were more successful in target Level 5 than Level 3 throughout (Level 3: 77.8 ± 14.4%, Level 5: 83.8 ± 12.1%; averaged across speeds and days, Figure 2A, right).

**Figure 2:**
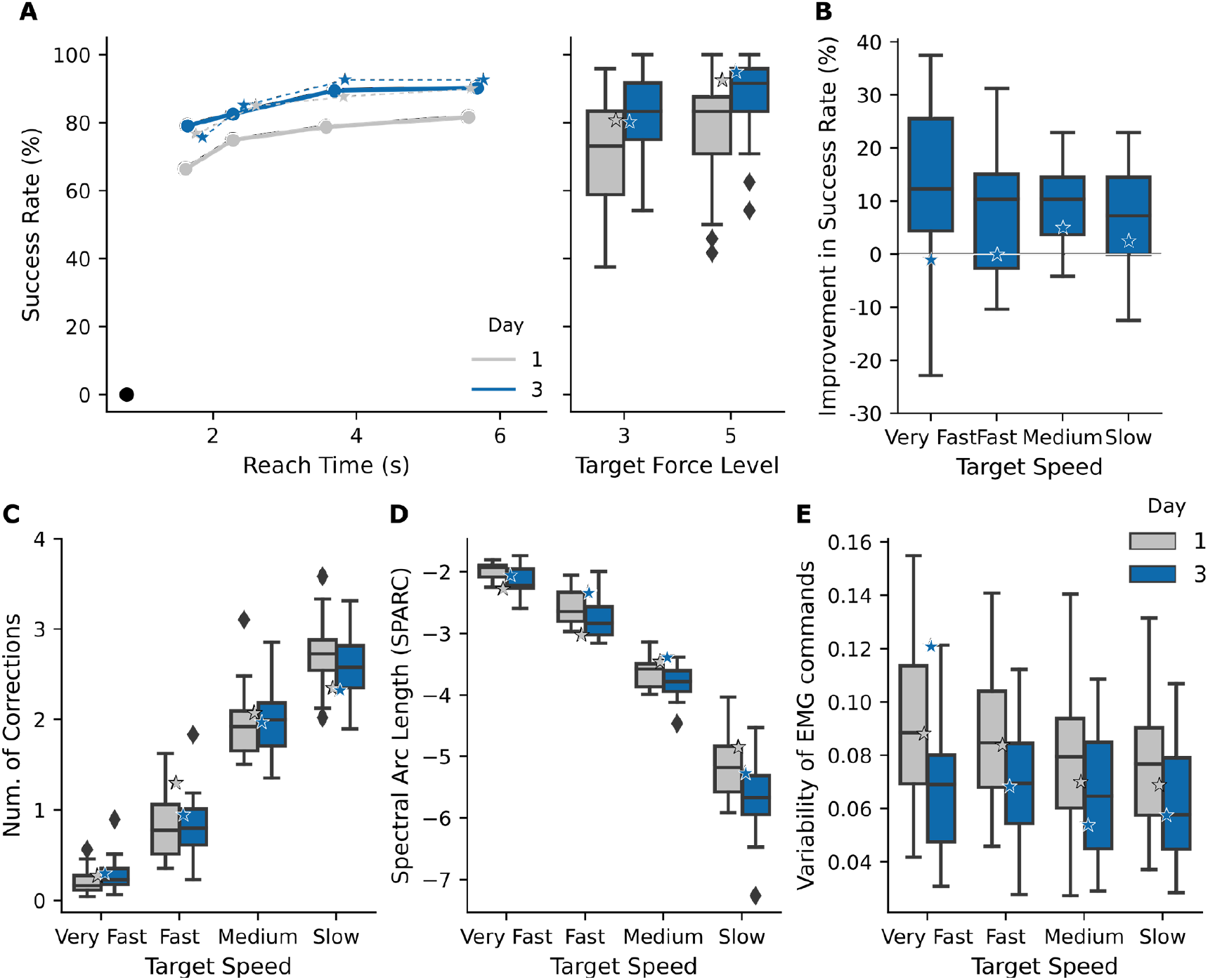
Learning induced changes in the speed-accuracy trade-off. (A) (left) Average SAF across able-bodied participants and the amputee is shown, on Day 1 and Day 3. Black circle on the bottom left indicates the x-intercept of SAF (success rate is necessarily 0% when reach time < 0.8 s). (right) Average success rates across speeds is higher for target force level 5. (B) Total improvement in performance (Day 1 to Day 3) is plotted per target speed. Notably, the improvement was similar across speeds. (C) Average number of corrections of force (p/trial) is plotted against target speeds. (D) Smoothness of generated EMG commands, measured by SPARC, decreased across days (greater values of SPARC implies smoother kinematics). (E) Trial-by-trial variability of EMG commands decreased across days for all speeds. Black diamonds indicate outliers, stars indicate amputee results.

Next, we analyzed how SAF changed between Day 1 and Day 3. Interestingly, while participants’ success rates improved across days, as explained above, there was no significant effect of target speed or target force on the observed improvement. That is, the increase in the success rate was similar across speeds though with large variability (Very Fast: 12.7 ± 15.6%, Fast: 7.6 ± 12.9%, Medium 9.8 ± 7.7%, and Slow 7.1 ± 9.7%, averaged across force levels, see Figure 2B), and across force levels (Level 3: 9.8 ± 8.8%, Level 5: 8.8 ± 8.7%, averaged across speeds). Therefore, the SAF curve effectively ‘shifted upwards’ across days consistently for all speeds, at both force levels.

Amputee results largely followed the trends of the able-bodied participants, however, possibly due to prior experience with similar experimental setups, her success rate improved only slightly from 84.8% on Day 1 to 86.3% on Day 3 (stars in Figure 2A, B).

### Behavioral Analyses

We then analyzed how participants’ behavior (planning and execution of EMG commands) changed across days. First, we investigated how participants’ control policies changed with training, by looking at the number of force corrections generated (Figure 2C, number of corrections are averaged over the two target forces to emphasize the effect of training (changes across days)). We found that the control policies were affected by the target speed (F(3, 45)=634.0, p<1e-10), and target force (F(1, 15)=843.7, p<1e-10). However, there was no significant effect of day, indicating that the overall strategy of how participants used the feedback remained consistent throughout the experiment. There was an interaction effect between target force and speed (F(3, 45)=115.8, p<1e-10), whereby the number of corrections for target level 3 decreased more gradually with speed, compared to level 5. The number of corrections decreased steadily with the target speed (post-hoc tests were significant for all pair, with each p-adj<1e-7), suggesting that at faster speeds, participants felt more pressed to adjust their muscle contractions prior to contacting the object, rather than using feedback post-contact to reach the required force. Finally, participants made fewer corrections to reach force level 3 than level 5 (p<1e-10).

Next, we analyzed how the smoothness and trial-by-trial variability of their movements were influenced by learning (Figure 2D, E, both measures are shown by averaging across target force levels, similar to Figure 2C). We found that there was a significant effect of all three independent variables – target speed, target force and day on both metrics^1^. Additionally, there was a significant interaction effect of target speed and target force on movement smoothness (F(3, 45)=69.8, p<1e-10), such that the smoothness of commands to reach target level 3 decreased more rapidly with speed, compared to level 5. Surprisingly, smoothness decreased from Day 1 to 3 (p-adj=6e-4), without any significant changes on consecutive days, while the smoothest trajectories were observed at the fastest speeds (post-hoc tests were significant for all pairs, with each p-adj<1e-8). On the other hand, variability improved (decreased) from Day 1 to 2 (p-adj=0.002) and then stagnated. Moreover, it was significantly different between the faster and slower speeds, with a significant difference between Fast and Medium (p-adj=0.046), but not between Fast and Very Fast or Medium and Slow. Finally, while participants produced smoother commands to reach force level 5 (compared to level 3 p=2e-6), variability of those commands was significantly larger than while reaching level 3 (p=9e-9). Behavioral results of the amputee participant also largely mirrored their able-bodied counterparts, except movement smoothness which improved across days (Figure 2C-E).

### Skill monitoring using ‘observational’ trials and SAF models

A power-law model of SAF with 2 parameters was fit to log-transformed performance data obtained by pooling across participants (see Section 2.8). Across the three days, the average R^2^ was 0.76 ± 0.09. Figure 3A displays the model fit in the log-log plane (and Figure 3B in true coordinates) where the data was pooled across subjects and demonstrates that the slope of the SAF remained similar across training (Day 1 to 3: -0.49, -0.57, -0.58), while the intercept decreased (Day 1 to 3: -1.18, -1.42, -1.63), which is in line with the conclusions of previous studies [14], [23]. Therefore, we proceeded to simplify the 2-parameter model by fixing the slope. To determine the parameter value of the slope that least affected the R^2^ of the model fits across the three days, we conducted a perturbation analysis where we fixed the intercept to its true value and varied the slope in the vicinity of the best-fit values obtained above. We found that when the slope was fixed at -0.55, we observed almost no loss in the goodness of fit across the three days (a decrease of 0.009 in R^2^). In comparison, fixing the intercept led to a decrease of at least 0.18 in R^2^. Therefore, we fixed the slope to be -0.55 for our 1-parameter model.

**Figure 3:**
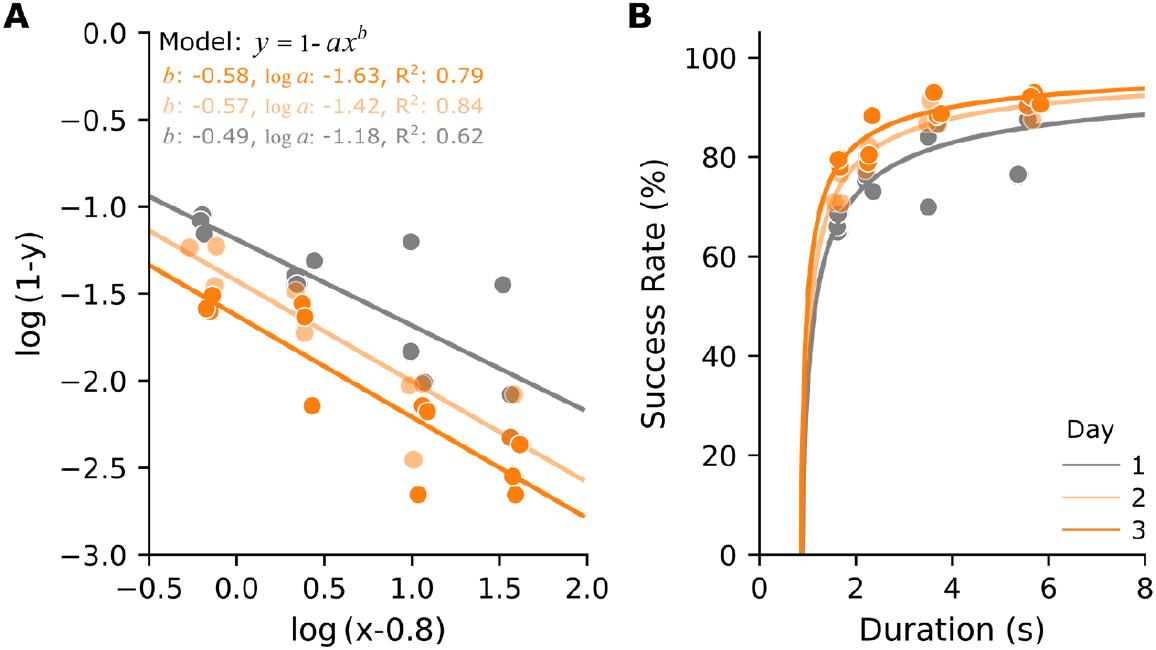
Power law models of SAF to infer skill. (A) Fitting the power-law model to (log-transformed) pooled participant data. Filled circles indicate measured success rates, solid lines indicate model fits. (B) Same data and models in (A), plotted in true coordinates.

Using this 1-parameter model, we proceeded to infer skill changes in an amputee case study. During the SAF trials, the participant’s success rate slightly improved (2.6% increase, averaged across speeds) from Day 1 to Day 3, which together with the improved trial-by-trial variability indicates a slight improvement in skill. Apart from the SAF experiment, where execution speed was controlled, the amputee participant performed ‘observational’ trials where the speed was unrestricted, on Day 2 and Day 3. During these trials, the average success rate increased from 89.1 ± 9.2% to 96.8 ± 3.1%, but with an accompanied increase in reach times (decreased speed) from 1.95 ± 0.3 s to 2.96 ± 0.27s, thereby making skill inference infeasible using just success rate and reach time data. We subsequently fit the 1-parameter SAF model, to analyze if there is a change in the intercept. We found that there is a decrease in the intercept from -2.14 to -3.02, indicating a trend towards improvement of skill. While the intercept values obtained were larger than those for the pooled participant data, this can be explained by the difference in average performance between the pooled participant (speed-restricted) trials and the amputee’s observational trials.

## Discussion

Speed-accuracy tradeoff functions (SAFs) provide a rigorous way to evaluate and understand motor skill and monitor its improvement across days. Here, we empirically measured how the SAF changed across three days for a prosthesis force control task where participants used a closed-loop interface based on proportional myoelectric control and EMG feedback. We found that the shape of the SAF curve remained intact, as participants’ success rate improved similarly across all speeds and forces. We then built parametric models of the measured SAFs to better characterize the changes in SAF and then discussed a method to monitor skill changes using observational data from an amputee participant.

### Learning induced changes in the SAF for an EMG Feedback-based Interface

In this study, we focused on an EMG feedback-based interface that has previously been shown to outperform other non-invasive feedback approaches to prosthesis force control [25], [31]. We elicited a wide range of success rates by enforcing task execution at different speeds using a time-band methodology, similar to the previous studies that investigated motor control of natural movements [16], [22]. We found a significant improvement in performance with training across all speeds. We expected that the largest gains in the SAF would occur toward faster speeds (Fast and Very Fast), due to the nature of EMG feedback which promotes predictive modulation of muscle contractions [11], [24]. However, conversely, we found that the SAF merely ‘shifted upwards’. Nevertheless, the shape of the SAF curve flattened up for the lower speeds (no difference between Medium and Slow), and it would be interesting to analyze whether prolonged training would eventually change the profile shape by flattening out the other side of the curve (i.e., similar performance across all speeds).

We also noticed that the shape of the SAF was similar across force targets. Interestingly, we found that the participants had greater success reaching target level 5 compared to 3, even at faster speeds. This could be attributed to the design of our piece-wise linear control mapping, which ensures that the variability at stronger contractions is compensated for by the width of the levels. Taken together, these outcomes are encouraging for the application of EMG feedback as they demonstrate that this interface affords a consistent and steady improvement in skill over the range of speeds and forces. In addition, most of the improvement happens quickly, after a single day of training (from Day 1 to Day 2). It remains to be tested if the lack of significant improvement between Days 2 and 3 means that this is the maximum performance or, more likely, that prolonged training would lead to further improvement but with more gradual increments.

Underlying these changes in performance, we observed that the behavioral outcome measures also exhibited interesting trends during training. First, there was no change, across days, in the strategy adopted by participants in each of the speed conditions, as shown by the number of corrections they used to achieve the target force. The fact that the number of corrections decreased with speed to less than 1 for Fast and Very Fast conditions shows that, when required, the participants successfully utilized predictive control afforded by EMG feedback and adjusted the contraction to a proper level even before the hand contacted the object. At the level of generated EMG commands, we observed that the generated myoelectric signals became less variable (or more similar to each other from trial to trial) across days, indicating that participants’ motor execution became more stereotypical, a hallmark of skilled behavior. However, counter intuitively, the commands also became less smooth. This could be due to the discretized nature of the task and feedback interface, which promotes participants to navigate from one level to the next, by ‘pausing’ in between to avoid overshooting, and thereby leading to less smooth trajectories as participants pause for short periods of time.

Interestingly, while the success rate of the amputee participant did not change substantially, both movement smoothness and variability improved in most cases (three out of four speeds). This underscores the relevance of measuring movement characteristics to better monitor user skill progression, as has also been highlighted in earlier studies [8], [9]. Moreover, it substantiates the utility of analyzing trial-by-trial variability of myoelectric commands as a marker of skill, analogous to reduced variability of end-point kinematics in natural movements [17], [18].

### Measuring and monitoring skill through SAF models

Speed and accuracy of movements are inextricably linked, and skill can be best understood as a combination of the two. SAFs therefore provide a quantitative framework through which to measure skill. In combination with previous results where SAF was introduced as a means of comparing and evaluating different closed-loop prosthesis interfaces (Force vs. EMG feedback) [22], here we observed that the SAFs so measured do not change shape across days, at least for the EMG feedback interface. Given this also matches the conclusions from the studies investigating the intact motor system, the constancy of tradeoff (both with respect to force levels and speeds) might be a more general phenomenon, as also expressed in [16]. This raises an interesting possibility – given that participants improve similarly across speeds, the tradeoff enabled by one interface may be different than another no matter the level of expertise of the user. For example, in [27], the benefits of EMG feedback with respect to force feedback were most expressed for medium speeds, and the present study implies that this might hold regardless of training. However, this still needs to be confirmed experimentally. Taken together, we expect that the SAF methodology used here is generally applicable across interfaces and force-matching tasks, with appropriate time scales.

Additionally, here we encapsulated the SAFs using a power law model, whereby the parameters of the model completely determined the SAF. Accordingly, empirically estimating the model parameters and investigating how they change over time gives us a powerful methodology to measure and monitor user skill progression. In line with previous efforts in the field of human motor control of natural movements, and human-computer interaction [14], [23], we observed that the exponent of the power law model (slope in log-transformed coordinates) remained relatively constant. We leveraged this knowledge to overcome one potential limitation of the SAF methodology – that participants need to repeat the same task at multiple speeds, requiring more time than the common practice of performing the task at a single self-chosen speed. This was demonstrated in the present study by determining the scale parameter (the intercept in log-transformed coordinates) from observational data while the slope parameter was set to an experimentally determined average value, to infer skill in a single amputee case study (in line with [14]), as opposed to merely comparing success rates across days. However, given the narrow variability in the observational data, it remains to be seen if a single execution speed sufficiently estimates the parameter of interest or if experimentally sampling more speeds might lead to better estimates. Due to this and other limitations outlined in the next section, the presented analysis and application of a single parameter model shall be regarded mostly as a proof-of-concept. Furthermore, while such a simplified SAF model perhaps enables skill inference and provides a good approximation, we believe that measuring the SAF and the underlying motor abilities, is substantially more informative.

### Limitations and future work

One limitation of the current study is the fact that we only conducted the observational experiment on the amputee subject. Moreover, we observed that they had very high levels of skill, already to begin with, having participated in similar experimental setups before, and therefore did not improve by much. Therefore, future studies need to verify the validity of the proposed model-based skill inference framework with a larger cohort of participants with measured SAF profiles in combination with observational data. However, we believe that the approach of modelling SAFs and incorporating that knowledge to infer skill represents a promising direction to promote rigorous skill inference, and for benchmarking efforts that characterize the performance afforded by different interfaces. The latter can also include the baseline condition, with no artificial sensory feedback, where prosthesis users can only rely on the incidental sources of information (visual and auditory cues). This would allow quantifying the state of the art in ‘open-loop’ user-prosthesis interfaces as well as reveal the benefits of added feedback.

## Conclusion

In this study, we empirically derived the changes in speed-accuracy tradeoffs (SAFs) during a 3-day experiment, as participants performed a functional prosthesis force-matching task. We found that success rates increased from Day 1 to Day 3, and that the improvement was similar across the target forces and target speeds. The training therefore improved performance without altering the initial shape of the SAF profile. We then modeled the empirically observed profiles to thoroughly characterize learning induced changes in SAF with EMG feedback. More generally, the study demonstrates that the SAF methodology can be successfully translated to prosthesis control for systematic and comprehensive monitoring of user skill.

## Acknowledgements

This work has been supported by the projects 8022-00243A (ROBIN) and 8022-00226B funded by the Independent Research Fund Denmark. We thank Jakob Nebeling Hedegaard for helpful discussions regarding SAF model fitting and comparison.

## Footnotes

1 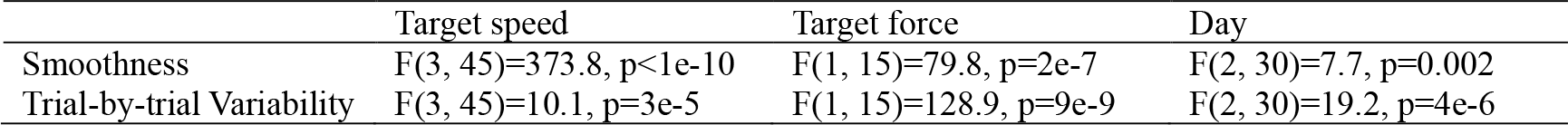

